# Ecological Dynamics of Pro-tumor and Anti-tumor Teams in the Tumor Microenvironment

**DOI:** 10.64898/2025.12.31.697148

**Authors:** Vaibhav Anand, Mohit Kumar Jolly, Herbert Levine

## Abstract

Tumor growth occurs within a complex tumor microenvironment (TME) composed of many interacting cell types. The immune cell types in TME tend to organize into two functional communities: a pro-tumor team and an anti-tumor team, each internally cooperative but mutually antagonistic forming a two-team ecosystem. Quantitatively predicting the ecological outcomes of such interactions remains challenging due to cellular diversity and interaction variability, and the exact dynamical regimes accessible to such a two-team ecosystem remain unknown. Here, we model tumor-immune interactions as a structured ecosystem with two competing teams using a generalized Lotka–Volterra framework and analyze it using the cavity method. We derive phase diagrams that delineate when these two communities coexist, when one dominates, and how these outcomes depend on intra-team cooperation, cross-team inhibition, and ecological heterogeneity. Our work provides a foundation for understanding tumor-immune dynamics from a community ecology perspective.

## 1 Introduction

Cancer is a systemic disease in which cells escape the normal constraints of growth and regulation. Among its classical hallmarks [1] are sustained proliferative signaling and the evasion of growth suppressors, which together render tumor cells quasi-autonomous entities capable of unchecked expansion. This growth unfolds within a highly complex environment composed of diverse cell types that collectively form the tumor microenvironment (TME). Non-malignant cells within this environment were once thought to play a passive role in tumorigenesis; it has now become evident that they actively shape tumor progression, collectively forming a structured ecological community [1–6]. Although the evolutionary dynamics of cancer cells have been studied extensively [7–11], how these dynamics unfold on shorter time scales through ecological interactions within the TME remains underexplored [12–16].

Within the tumor microenvironment(TME), several kinds of ecological interactions occur among cancer cells, immune cells, and stromal components. Cancer cells are subject to immune surveillance and also compete with host cells (and other resident cells) for limited resources such as oxygen, nutrients, and space. Immune and stromal cell types can be broadly grouped into two functional communities: an anti-tumor group arising from the adaptive and innate immune responses that target cancer cells, and a pro-tumor group typically driven by tumor-derived signals that recruit and reprogram inflammatory or immunosuppressive cells to secrete growth factors that support tumor expansion [3, 5, 18]. For instance, macrophages exhibit two phenotypes, M1 (anti-tumor) and M2 (pro-tumor) [19]. Similarly, tumor-associated neutrophils exist in two distinct phenotypes, N1 (anti-tumor) and N2 (pro-tumor) [20]. Moreover, phenotypic plasticity has been observed between effector T cells (Teffs; anti-tumor) and regulatory T cells (Tregs; pro-tumor) [21, 22]. Cell types within each group tend to cooperate with others in the same group and antagonize those in the opposing group, through mechanisms such as recruitment and/or activation of team players (see Fig. 1). This configuration implies an ecological system composed of two competing teams [17], but the precise dynamical regimes accessible to such a system remain unknown.

**Figure 1:**
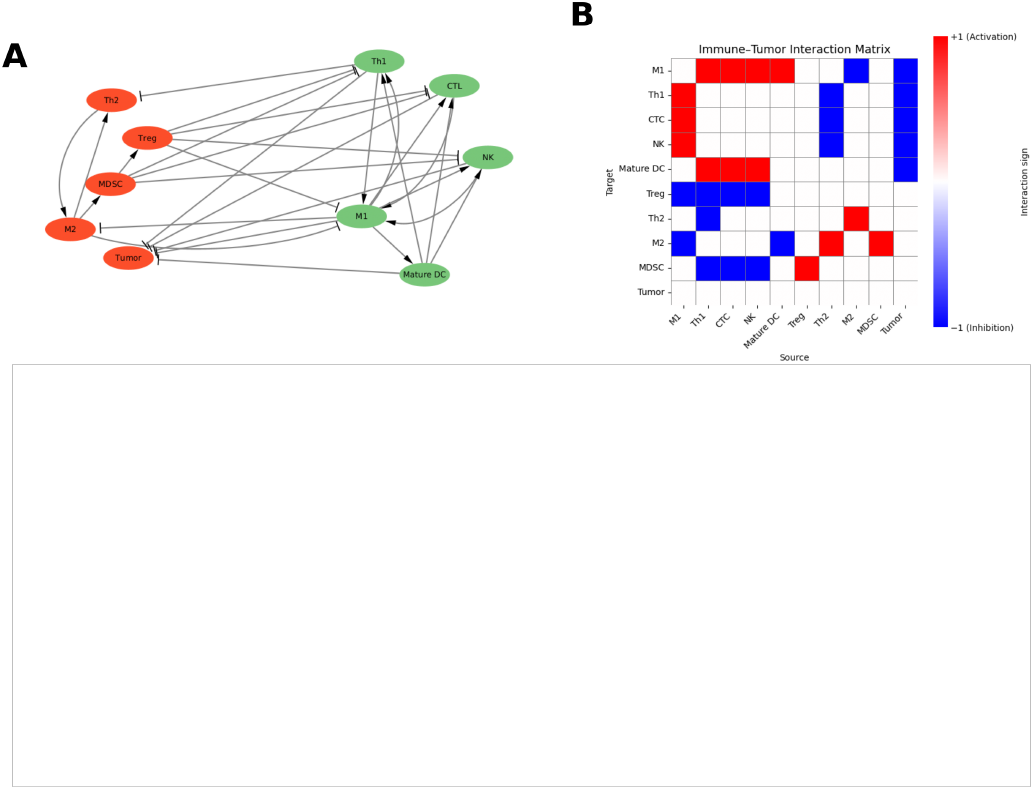
Teams in the tumor microenvironment. (A) Network representation of interactions among major cell types in the TME, adapted from [17], reveals two distinct communities: the anti-tumor team (green) and the pro-tumor team (red). (B) Adjacency matrix illustrating activating and inhibitory interactions between cell types.

A major challenge in modeling these ecosystems lies in their high dimensionality, which makes quantitative analysis difficult. Yet studies of ecological networks suggest that this apparent complexity often masks simpler underlying structure. Many real ecosystems can be decomposed into only a few functional modules [23], and their effective dimensionality is frequently much lower than expected [24]. Bipartite mutualistic networks provide a clear example of such structured ecological architecture, and have been characterized in detail [25, 26]. Motivated by these insights, we investigate ecosystems composed of two subcommunities that cooperate internally while antagonizing each other.

The generalized Lotka–Volterra (GLV) equations offer a natural mathematical framework for analyzing species rich ecosystems and their structures [27, 28]. These models have been widely studied using the cavity method, which enables the determination of equilibrium statistics in the limit of large system size, by treating species-level fluctuations as small and interaction parameters as drawn from statistical ensembles [29–33]. Predictions from such models have shown strong agreement with more detailed variants [34].

Here, we study a structured two-team ecosystem driven by GLV dynamics, with interaction parameters drawn from Gaussian ensembles whose variances encode ecological heterogeneity. Using the cavity method, we derive equilibrium statistics, including survival fractions and mean abundances for both teams. By tuning the balance between intra-team cooperation and inter-team inhibition, we uncover transitions between mutual coexistence, competitive exclusion of one team, and unbounded proliferation of one or both the teams. We further show that heterogeneity in interaction strengths within and across teams shapes these transitions in distinct ways. Taken together, these results provide a statistical physics based ecological framework for understanding the collective dynamics between pro-tumor and anti-tumor factors in the tumor microenvironment.

## 2 Results

### 2.1 Effect of heterogeneity on ecosystems with teams

As already mentioned, we model pro-tumor and anti-tumor cell types as two interacting teams of species, whose dynamics are governed by the generalized Lotka–Volterra (GLV) equations. In the simplest unstructured community setting, the abundance *N*_*i*_ of species *i* evolves according to

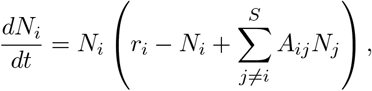

where *N*_*i*_ denotes the normalized abundance of species *i* (scaled by its carrying capacity) and *S* is the total number of species in the community. Here, *A*_*ij*_ represents the effect of species *j* on *i, A*_*ij*_ *<* 0 indicates inhibition of *i* by *j*, and *A*_*ij*_ *>* 0 indicates activation of *i* from *j*. The intrinsic growth rates are drawn from a normal distribution *r*_*i*_ ∼ 𝒩 (*µ*_*r*_, *ζ*).

In the cavity method framework, the interaction coefficients are defined as

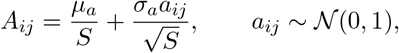

where ⟨*a*_*ij*_, *a*_*ji*_⟩ = *γ*. For simplicity, we will focus on the case *γ* = 0. The 1*/S* and 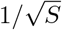 scaling of the mean and variance of *A*_*ij*_ ensures a well-defined thermodynamic limit as the number of species *S* becomes large. Specifically, the 1*/S* factor normalizes the cumulative contribution of all interactions so that the mean interaction effect on each species remains 𝒪 (1), while the 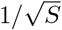 factor guaranties that the fluctuations arising from random interactions also remain finite. This scaling allows the macroscopic properties of the system—such as species survival fractions, mean abundances, and stability—to converge to deterministic values as *S* → ∞.

We can then derive precise statistics of a random community (*ϕ*, ⟨*N* ⟩, ⟨*N*^2^⟩ for a given *µ* and *σ* defining the dynamical ensemble. In unstructured random communities, when the heterogeneity (*σ*) can is above a certain threshold, we enter a disordered regime of multiple attractors and chaotic dynamics (Fig. 2A, 2C) [29][30]. Otherwise, we have steady-state co-existence of various species, as can be seem by simulations of specific ecosystem realizations in this parameter range (Fig. 2B).

**Figure 2:**
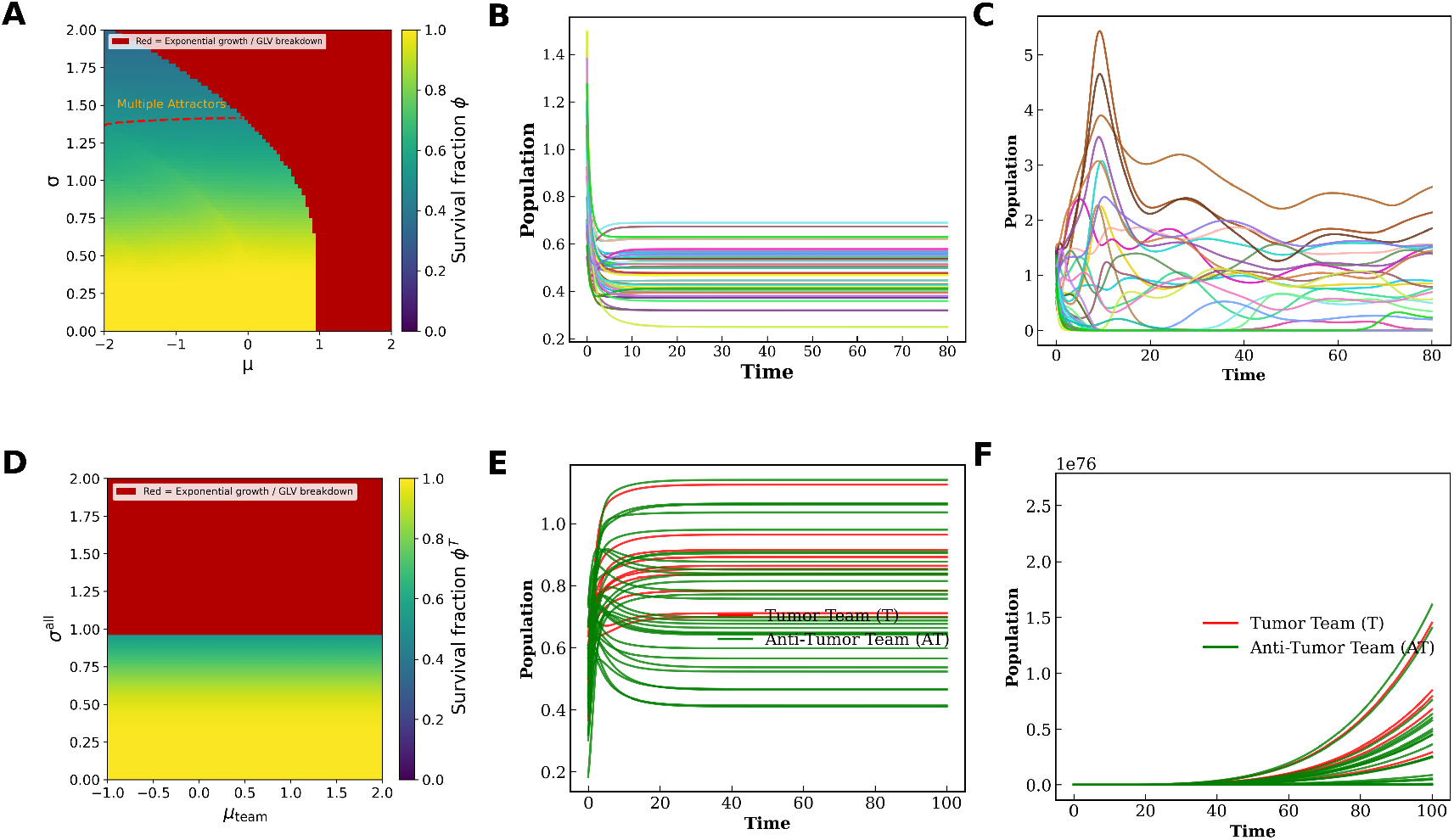
High heterogeneity leads to extinctions in ecosystems with teams. (A) Survival fraction for an unstructured random ecosystem in the (*σ, µ*) plane (*γ* = 0). The dashed red line marks the onset of the multiple-attractor phase; red shading indicates cavity-method non-convergence. (B) Dynamics of a 20-species GLV system sampled from the unique-fixed-point phase (*µ* = −1, *σ* = 0.2) and (*µ*_*r*_ = 1, *ζ* = 0) in the unstructured random ecosystem showing stable convergence. (C) The same system sampled at high heterogeneity (*µ* = −1, *σ* = 1.6), showing unbounded trajectories. (D) Phase diagram for tumor team in a two team structured ecosystem in (*µ*^team^, *σ*_all_), with *µ*^*T*−*T*^ = *µ*^*A*−*A*^ = *µ*^team^ and 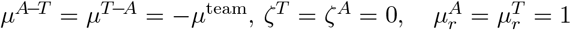. Increasing *µ*^team^ strengthens both intra-team cooperation and inter-team antagonism. (E) GLV simulation of two 20-species teams at *σ*_all_ = 0.2, *µ*^*T*−*T*^ = *µ*^*A*−*A*^ = 0.5, *µ*^*T*−*A*^ = *µ*^*A*−*T*^ = −1, showing stable coexistence. (F) The same system at *σ*_all_ = 1.6, showing unbounded growth.

We now generalize the method to a structured ecosystem with two teams, a pro-tumor team and an anti-tumor team. the dynamics of the *i*^th^ species belonging to tumor team (tumor team denoted by T and anti-tumor team denoted by A) are given by

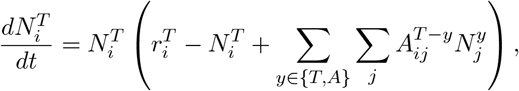

and that of *i*^th^ species belonging to anti-tumor team are given by

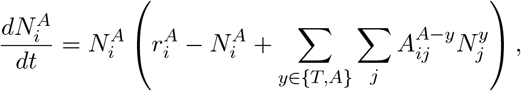

where 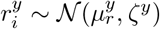(for *y* = {*A, T*}) represents the group-specific growth rate distribution. Each group *T/A* consists of *S*_*T/A*_ species, and the total number of species in the ecosystem is *S* = *S*_*T*_ +*S*_*A*_. The interaction matrix between species belonging to groups *T* and *A* is defined as

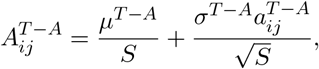

with 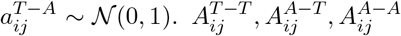 are defined similarly. We denote the tumor team by T and anti-tumor team by A and therefore *µ*^*T* −*T*^, *µ*^*A*−*A*^, *µ*^*A*−*T*^, *µ*^*T* −*A*^ are the means for the interactions within the pro-tumor team, within the anti-tumor team, on the anti-tumor team from the pro-tumor team and from the anti-tumor team on the pro-tumor team respectively.

Extending the cavity method to this case (see Appendix), we are able to solve for survival fraction *ϕ*^*T/A*^, first moment/mean population ⟨*N*^*T/A*^⟩ and second moment ⟨(*N*^*T/A*^)^2^⟩ for each group given the interaction means *µ*^*T* −*T*^, *µ*^*A*−*A*^, *µ*^*T* −*A*^, *µ*^*A*−*T*^, and standard deviations *σ*^*T* −*T*^, *σ*^*A*−*A*^*σ*^*A*−*T*^ *σ*^*T* −*A*^, as well as the growth rate standard deviation *ζ*^*T*^, *ζ*^*A*^. Each team occupies a fraction 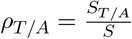 of the total community, allowing us to analyze inter-team and intra-team interactions in a tractable manner.

To begin to contrast the pro-tumor/anti-tumor team ecosystem with an unstructured random community, we define a composite parameter *µ*_*teams*_ which captures the balance between cooperation (within members belonging to same team) and inhibition (among members belonging to separate teams), we define a composite parameter,

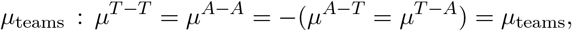

and define *σ*^*all*^ = *σ*^*T* −*T*^ = *σ*^*A*−*T*^, *σ*^*T* −*A*^, *σ*^*A*−*A*^ We have a two-team ecosystem when *µ*_teams_ *>* 0, and larger values indicate stronger competition between the two teams with stronger cooperation within the teams. On the other hand, *µ*_teams_ *<* 0 denotes the case where there is antagonism within the teams but cooperation between species in different teams. This may arise in a system with competition among related species for the benefits arising from mutualistic interactions with other parts of the ecosystem.

More specifically, in fully random ecosystems, the sign of the mean interaction strength sets the overall ecological “mode”: *µ >* 0 corresponds to net cooperation, while *µ <* 0 corresponds to net competition. In our two-team ecosystem with positive *µ*_teams_, however, the situation is more nuanced—cooperative interactions occur *within* each community, while competitive interactions act *between* them. Despite this structural difference, both random ecosystems and the two-team ecosystem display a common trend: increasing interaction heterogeneity drives species extinctions.

As we have seen, in random ecosystems once the variance of interactions exceeds a critical threshold, the system transitions from a regime with stable abundances to a multiple-attractor phase with fluctuating abundances. Simulations in the former regime produce clean, stable convergence for a 20-species community (Fig. 2B), whereas simulations in the multiple-attractor regime show highly variable non-steady outcomes (Fig. 2C). The red region in Fig. 2A marks where the cavity equations do not converge, corresponding to unbounded growth. This typically occurs when cooperation dominates (*µ >* 0), but sufficiently high heterogeneity can also destabilize systems that are, on average, competitive (*µ <* 0). Importantly, the amount of heterogeneity required for this blow-up increases as the baseline competition becomes stronger.

The two-team ecosystem behaves differently. Here, “competition balances competition”, namely strong antagonistic interactions between teams stabilize the system, and convergence persists even at high values of *µ*_team_. We find that cavity-solver breakdown in the two-team model occurs only at higher heterogeneity (approximately *σ >* 1.0). For *σ*_all_ *<* 1, both teams robustly coexist, consistent with direct simulations of finite ecosystems with 20 species per team. Unlike the random ecosystem, in the structured ecosystem, due to teams structure, we reach GLV breakdown before the multiple attractor regime which is consistent with simulations (Fig. 2D).

### 2.2 An ecosystem with two competing teams results in distinct regimes of co-existence, exclusion and unbounded proliferation

So far, we have examined the role of a single parameter *µ*_teams_ that enforced identical intra–community cooperation. However, in an ecosystem, the two teams could have different strengths of coopera-tion and competition. Here, we generalize this analysis by independently varying *µ*^*T* −*T*^ and *µ*^*A*−*A*^. Throughout this section, we fix *σ*^all^ = 0.2 and assume that both communities are of equal size (*ρ*^*T*^ = *ρ*^*A*^ = 0.5). The inter–team interactions are taken to be inhibitory, with *µ*^*T* −*A*^ = *µ*^*A*−*T*^ = −1, reflecting the antagonistic nature of the tumor–versus–anti–tumor interplay.

For each pair (*µ*^*T* −*T*^, *µ*^*A*−*A*^), we solve the cavity equations self-consistently to obtain the survival fractions and mean population levels of the two communities, namely *ϕ*^*T*^, *ϕ*^*A*^, ⟨*N*^*T*^ ⟩, and ⟨*N*^*A*^⟩. A two team ecosystem with intra team cooperation and inter team antagonism exists when *µ*^*T*^, *µ*^*A*^ *>* 0

As seen previously in Fig. 2E, species from both teams coexist with comparable population sizes when the teams are identical with *µ*^*T* −*T*^ = *µ*^*A*−*A*^ = 0.5. In the more general (*µ*^*T* −*T*^, *µ*^*A*−*A*^) plane, we observe coexistence between the tumor and anti–tumor communities whenever *µ*^*T* −*T*^, *µ*^*A*−*A*^ ≤ 1. This regime includes cases with inhibitory intra–team interactions (*µ*^*T* −*T*^, *µ*^*A*−*A*^ ≤ 0), yet both teams maintain nonzero survival fractions (Fig. 3A,3B).

**Figure 3:**
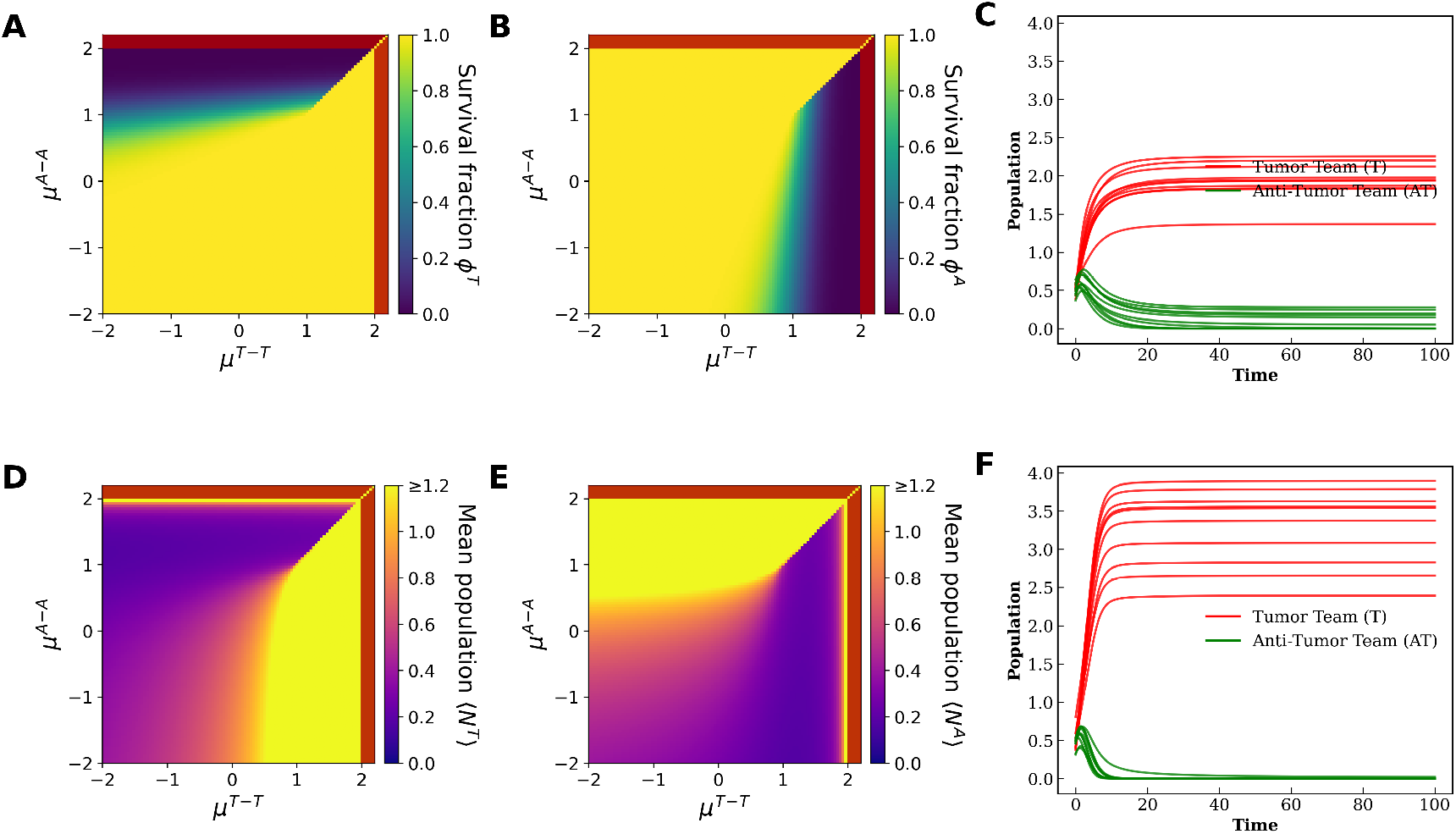
Phase diagrams of an ecosystem with two teams. Self-consistent equations for tumor and anti-tumor teams solved in the (*µ*^*T* −*T*^, *µ*^*A*−*A*^) space under inhibitory interactions (*µ*^*T* −*A*^ = *µ*^*A*−*T*^ = −1) with heterogeneity (*σ*^*T* −*T*^ = *σ*^*A*−*A*^ = *σ*^*A*−*T*^ = *σ*^*T* −*A*^ = 0.2, *ζ* = 0). Team fractions are fixed at *ρ*_*T*_ = *ρ*_*A*_ = 0.5. Red region indicates non convergence and GLV breakdown. (A) Tumor team survival fraction *ϕ*^*T*^, (B) Anti-tumor team survival fraction *ϕ*^*A*^, (C) GLV simulated with 20 species in each team where the intra team entries in the interaction matrix were sampled from the regime with *µ*^*T* −*T*^ = *µ*^*A*−*A*^ = 1.2 with *µ*^*T* −*A*^ = *µ*^*A*−*T*^ = −1 and *σ*^*all*^ = 0.2 (D) Tumor team mean population ⟨*N*^*T*^ ⟩, (E) Anti-tumor team mean population ⟨*N*^*A*^⟩, and (F) GLV simulated with 20 species in each team where the intra team entries in the interaction matrix were sampled from the regime with *µ*^*T* −*T*^ = *µ*^*A*−*A*^ = 1.2 with *µ*^*T* −*A*^ = *µ*^*A*−*T*^ = −1 and *σ*^*all*^ = 0.2

As cooperation within one of the communities becomes sufficiently strong, the system transitions into a regime where the weaker team begins to lose a significant fraction of its species. Eventually, the community with stronger cooperation dominates, maintaining a larger mean population ⟨*N* ⟩, while the other team approaches partial (Fig. 3C) or complete exclusion (Fig. 3F). The location of this coexistence–to–exclusion transition depends sensitively on the inhibitory strength between the two teams: stronger inter–team inhibition shifts the transition to higher levels of intra–team cooperation. We note that the yellow regions in the ⟨*N*^*T/A*^⟩ (Fig. 3D,3E) plots correspond to mean species abundances exceeding ∼ 1.2. Since abundances in our model are normalized by each species’ intrinsic carrying capacity, values above 1 reflect emergent, collective amplification driven by cooperative interactions within the team.

When both intra–team cooperative parameters satisfy *µ*^*T* −*T*^, *µ*^*A*−*A*^ ≥ 1, the system enters a strongly competitive regime. Here, the team with the strongest cooperative strength “wins” outright, maintaining *ϕ*^*T/A*^ = 1 and ⟨*N*^*T/A*^⟩ ≥ 1, while the weaker team undergoes complete extinction (*ϕ*^*T/A*^ = 0, ⟨*N*^*T/A*^⟩ = 0) (Fig. 3A, 3B, 3D, 3E). In this region, we observe abrupt transitions in both the survival fraction and the mean populations of the two teams. Finally, when *µ*^*team*^ ≈ 2 we enter a high cooperation regime that leads to the breakdown of the GLVs and leads to unbounded proliferation. This region is denoted by red in the plots (Fig. 3A, 3B, 3D, 3E). The various regimes are summarized in Table 1.

**Table 1:** Survival fractions of tumor and anti-tumor teams for different ranges of interaction mean values.

| $(\mu^{T-T}, \mu^{A-A})$ | $\phi^T$ (Tumor team) | $\phi^A$ (Anti-tumor team) | **Regime** |
| --- | --- | --- | --- |
| $(\mu^{T-T} \leq 1, \mu^{A-A} \leq 1)$ | $\phi^T \geq 0$ | $\phi^A \geq 0$ | Coexistence |
| $(\mu^{T-T} \geq 1, \mu^{A-A} \leq 1)$ | $\phi^T \approx 1$ | $\phi^A < 1$ | Tumor team dominates |
| $(\mu^{T-T} \leq 1, \mu^{A-A} \geq 1)$ | $\phi^T < 1$ | $\phi^A \approx 1$ | Anti-tumor team dominates |
| $(\mu^{T-T} \geq \mu^{A-A} \geq 1)$ | $\phi^T \approx 1$ | $\phi^A \approx 0$ | Anti-tumor team exclusion |
| $(\mu^{A-A} \geq \mu^{T-T} \geq 1)$ | $\phi^T \leq 1$ | $\phi^A \approx 0$ | Tumor team exclusion |
| $(\mu^{A-A} \geq 2, \mu^{T-T} \geq 2)$ | $\phi^T = 1$ | $\phi^A = 1$ | GLV breakdown/Non-convergence |

### 2.3 Simulations recapitulate analytical results

To check in more detail the accuracy of the cavity approach, we compare the equilibrium statistics obtained from the cavity method with those from direct numerical simulations of a two-team ecosystem with *n*_*T*_ = *n*_*A*_ = 10 species per team. The simulations were performed using (*µ*^*T* −*T*^, *µ*^*A*−*A*^) as the control parameters, with fixed inter-team means *µ*^*T* −*A*^ = *µ*^*A*−*T*^ = −1, and heterogeneity in parameters (*σ*^*T* −*T*^ = *σ*^*A*−*A*^ = *σ*^*A*−*T*^ = *σ*^*T* −*A*^ = 0.2, *ζ* = 0). The phase diagrams of survival fractions and mean abundances for both teams reproduce the regimes predicted by the cavity method. In particular, reduced survival fractions for both teams are observed when *µ*^*T* −*T*^ and *µ*^*A*−*A*^ exceed unity and are comparable in magnitude (*µ*^*T* −*T*^ ≈ *µ*^*A*−*A*^), indicating strong inter-team competition (Fig.4 A–D).

Quantitatively, the mean team abundances obtained from simulations closely match those predicted by the cavity method across a broad parameter range (Fig.4 E–F). The largest deviations occur for *µ*^*T* −*T*^ = 1.5, corresponding to the strongly cooperative regime, where we observe sharp transitions in both pro-tumor and anti-tumor mean populations and survival fractions as *µ*^*A*−*A*^ decreases below 1.5 (Fig.4 E, F). According to the cavity method, the tumor-team survival fraction *ϕ*^*T*^ abruptly increases from 0 to 1, while the anti-tumor survival fraction *ϕ*^*A*^ drops from 1 to 0 as the system enters the cooperative regime. These shifts are mirrored by corresponding transitions in the mean abundance levels. Even in this case, the discrepancy between the two approaches remains primarily quantitative—the cavity and simulation results exhibit strong qualitative agreement. For instance, the mean pro-tumor team abundance is ⟨*N*^*T*^ ⟩ ≈ 3 in simulations and ≈ 4 in the cavity prediction, both exceeding the carrying capacity (*K* = 1), consistent with a cooperative amplification of equilibrium abundance.

### 2.4 Community fraction and extinction boundaries

We next looked at the effect of the relative size of the two teams on dynamics. To do this, we varied the fraction of pro-tumor team (*ρ*^*T*^) and the strength of the anti-tumor antagonism on the pro-tumor team (*µ*^*T* −*A*^) to examine the effect of antagonism from anti-tumor team on the size of pro-tumor team and how it affects survival. As always, negative *µ*^*T* −*A*^ values represent inhibition, while positive values correspond to activation from the anti-tumor team. As expected, cooperative influence (*µ*^*T* −*A*^ *>* 0) leads to robust survival of the pro-tumor team, with survival fraction *ϕ*^*T*^ ≈ 1 and mean population ⟨*N*^*T*^ ⟩ *>* 1 (Fig. 4A,E).

**Figure 4:**
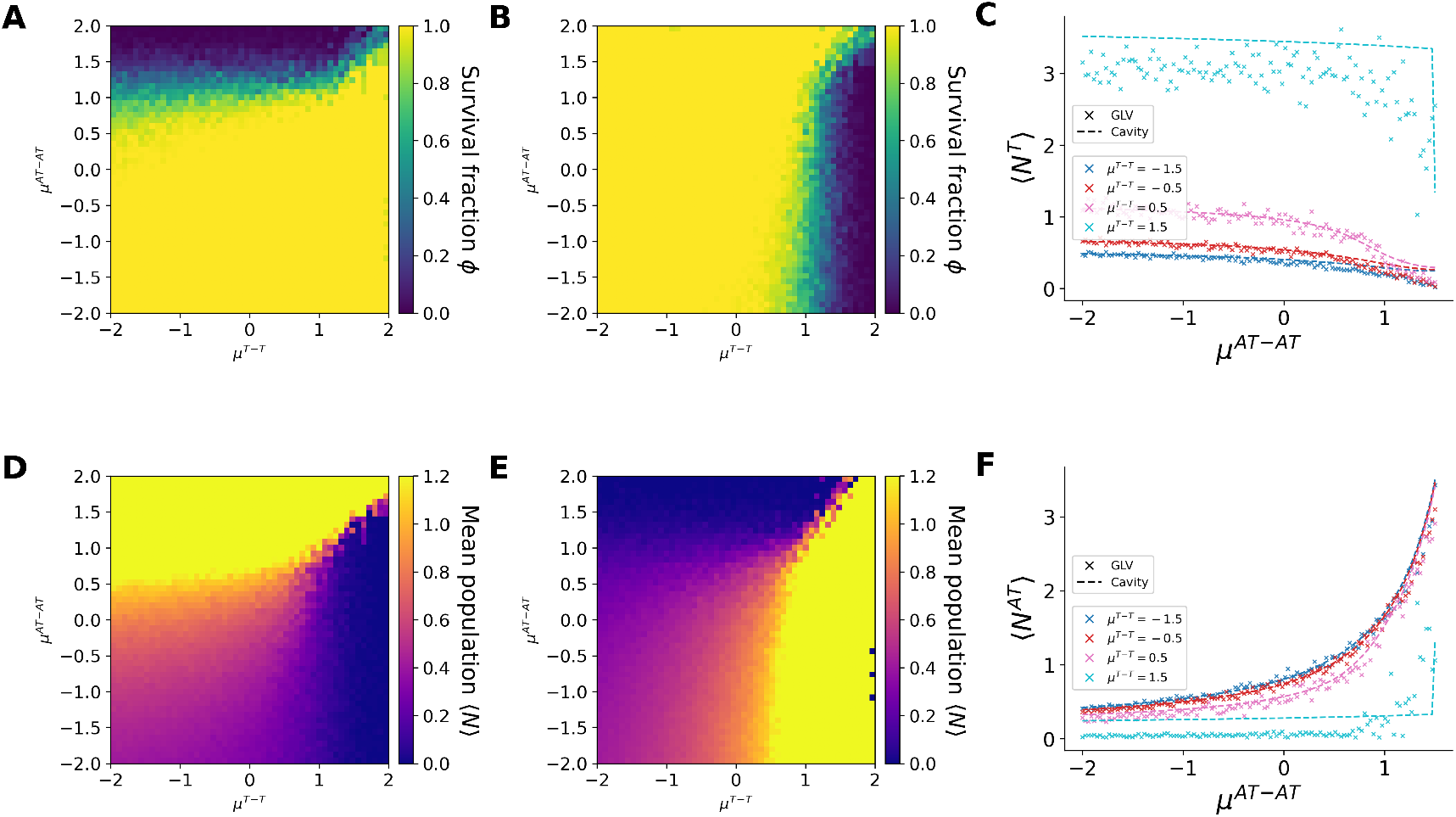
The cavity method accurately reproduces equilibrium population statistics. (A) Tumor-team survival fraction *ϕ*^*T*^ in *µ*^*T* −*T*^ and *µ*^*A*−*A*^ space. Each pixel represents the average over 10 stochastic replicates of GLV simulations with *n*_*T*_ = *n*_*A*_ = 10, total population size *S* = 20, and carrying capacities *r*_*i*_ = 1. (B) Anti-tumor survival fraction *ϕ*^*A*^ over the same parameter space. (D) Mean tumor-team abundance ⟨*N*^*T*^⟩ in the *µ*^*T* −*T*^ –*µ*^*A*−*A*^ space, normalized by the carrying capacity (*K* = 1). (E) Mean anti-tumor abundance ⟨*N*^*A*^⟩ . (C) Mean tumor abundance ⟨*N*^*T*^ ⟩ as a function of *µ*^*A*−*A*^ for fixed *µ*^*T* −*T*^ values. Solid (dashed) lines denote cavity (GLV) results, with per-curve *R*^2^ between generalized Lotka–Volterra (GLV) simulations and cavity-method predictions for the tumor mean abundance ⟨*N*_*T*_ ⟩, reported for different values of the intra-tumor interaction strength *µ*^*T*−*T*^ : *µ*^*T*−*T*^ = −1.5 (*R*^2^ = 0.673), −0.5 (0.836), 0.5 (0.919), and 1.5 (−1.842).. (F) Mean anti-tumor abundance ⟨*N*^*A*^⟩ for the same parameter sweep, increasing with *µ*^*A*−*A*^ as ⟨*N*^*T*^ ⟩ declines (*R*^2^) between generalized Lotka–Volterra (GLV) simulations and cavity-method predictions for the anti-tumor mean abundance ⟨*N*_*AT*_ ⟩, reported for different values of the intra-tumor interaction strength *µ*^*T*−*T*^ : *µ*^*T*−*T*^ = −1.5 (*R*^2^ = 0.985), −0.5 (0.980), 0.5 (0.964), and 1.5 (0.002).

Because *ρ*^*T*^ +*ρ*^*A*^ = 1, smaller tumor fractions (*ρ*^*T*^ *<* 0.6) shift dominance towards the anti-tumor team. Under inhibitory interactions (*µ*^*T* −*A*^ ∈ [−2, −1]), the pro-tumor community collapses, while moderate inhibition (*µ*^*T* −*A*^ *>* −1) allows the coexistence even for smaller pro-tumor team fractions (Fig. 5B, 5F). More generally, in the (*µ*^*T* −*T*^, *ρ*^*T*^) space, strong intra-team cooperation (*µ*^*T* −*T*^ ≥ 1) supports pro-tumor team survival even when *ρ*^*T*^ ≲ 0.5, though the pro-tumor team gets excluded when *ρ*^*T*^ *<* 0.4 even at high cooperation (*µ*^*T* −*T*^ = 2).

**Figure 5:**
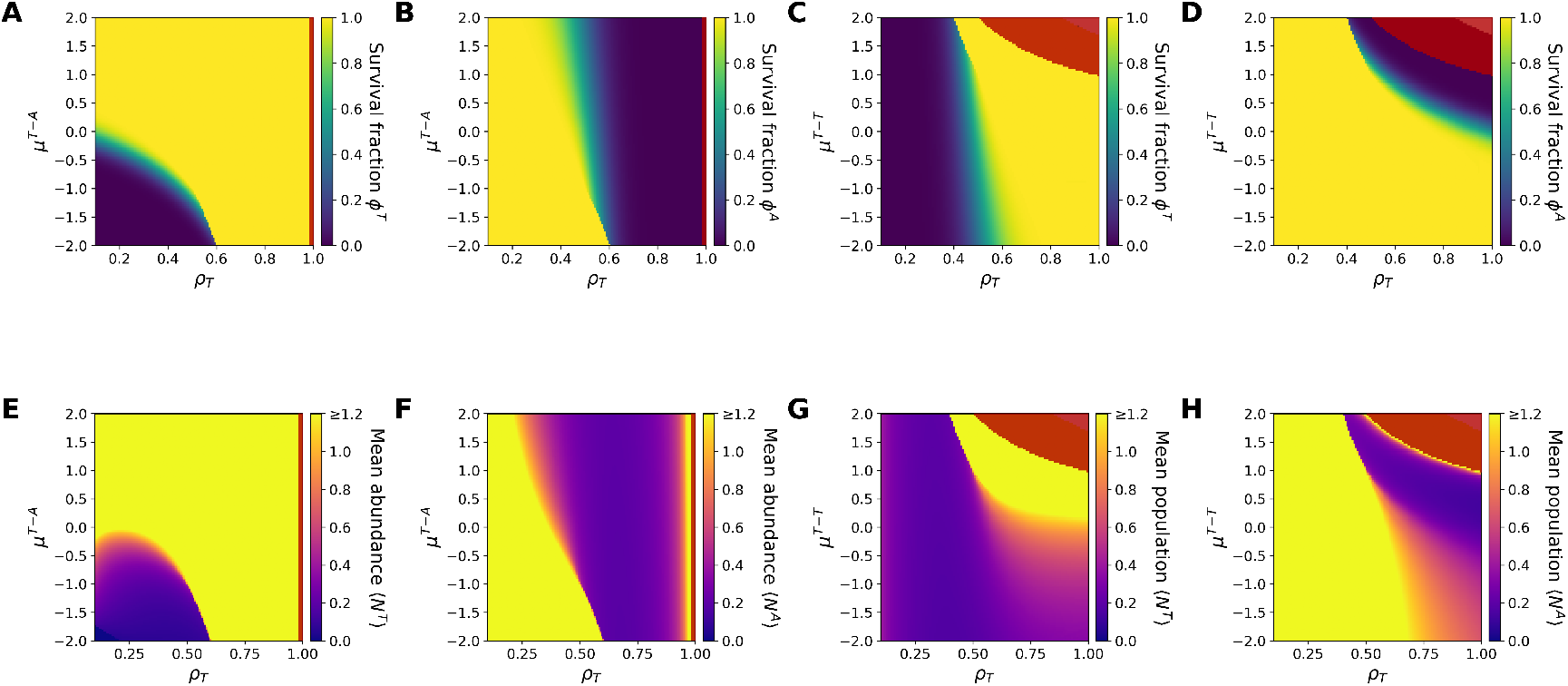
Community size determines persistence and extinction regimes. Self-consistent equations for tumor and anti-tumor teams with inhibitory interactions (*µ*^*T* −*A*^ = *µ*^*A*−*T*^ = −1) and no heterogeneity in parameters (*σ*^*T* −*T*^ = *σ*^*A*−*A*^ = *σ*^*A*−*T*^ = *σ*^*T* −*A*^ = 0.2, *ζ* = 0). Solved for tumor team survival fraction (*ϕ*^*T*^) and anti-tumor (*ϕ*^*A*^) team survival fractions in *µ*^*T* −*A*^ ∈ (−2, 2) and *ρ*^*T*^ ∈ (0.1, 1) space for A) Tumor team and B) Anti-tumor team. (C–D) Corresponding survival fractions plotted against *ρ*^*T*^ and intra-team interaction strength *µ*^*T* −*T*^ ∈ (−2, 2). (E–F) populations ⟨*N*^*T*^ ⟩ and ⟨*N*^*A*^⟩ in the (*ρ*^*T*^, *µ*^*T* −*A*^) space. (G–H) Mean populations in the (*ρ*^*T*^, *µ*^*T* −*T*^) space.

When inter-team inhibition is fixed (*µ*^*T* −*A*^ = *µ*^*A*−*T*^ = −1), the pro-tumor community survives only above *µ*^*T* −*T*^ ≥ 1 and at moderate *ρ*^*T*^ . Below *ρ*^*T*^ *<* 0.4, extinction occurs regardless of the cooperation strength. The viable region expands slightly when the anti-tumor team increases its internal cooperation, but remains narrow. At large tumor fractions (*ρ*^*T*^ *>* 0.5) and high self-activation (*µ*^*T* −*T*^ ≫ 1), the system exhibits explosive growth and breakdown of steady-state solutions, marking unbounded proliferation for the pro-tumor team (red regions in Fig. 5C, 5D, 5G, 5H).

### 2.5 Intermediate heterogeneity promotes coexistence

After investigating how different configurations of team-level mean interaction strengths shape ecosystem outcomes, we next examine the role of intra–team and inter–team heterogeneity. As shown in Fig. 2D, high overall heterogeneity *σ*^all^ leads to widespread extinctions in symmetric teams with equal cooperation, equal size, and equal competition. Extending this analysis, we computed survival fractions and mean abundances for asymmetric cooperation across the (*µ*^*T* −*T*^, *µ*^*A*−*A*^) plane.

To understand how heterogeneity alters the regimes identified in Section 2.2, we explored a range of *σ*^all^ values. While very high heterogeneity (*σ*^all^ ≥ 1) drives the system into an unbounded-growth regime, moderate heterogeneity (*σ*^all^ = 0.5) eliminates the sharp coexistence–exclusion transitions observed in Fig. 3A–E. In this intermediate regime, we also observe the disappearance of regions where either team maintains *ϕ* = 1; instead, both the pro-tumor and anti-tumor teams lose a finite fraction of their species throughout the phase space (Fig. 6A).

**Figure 6:**
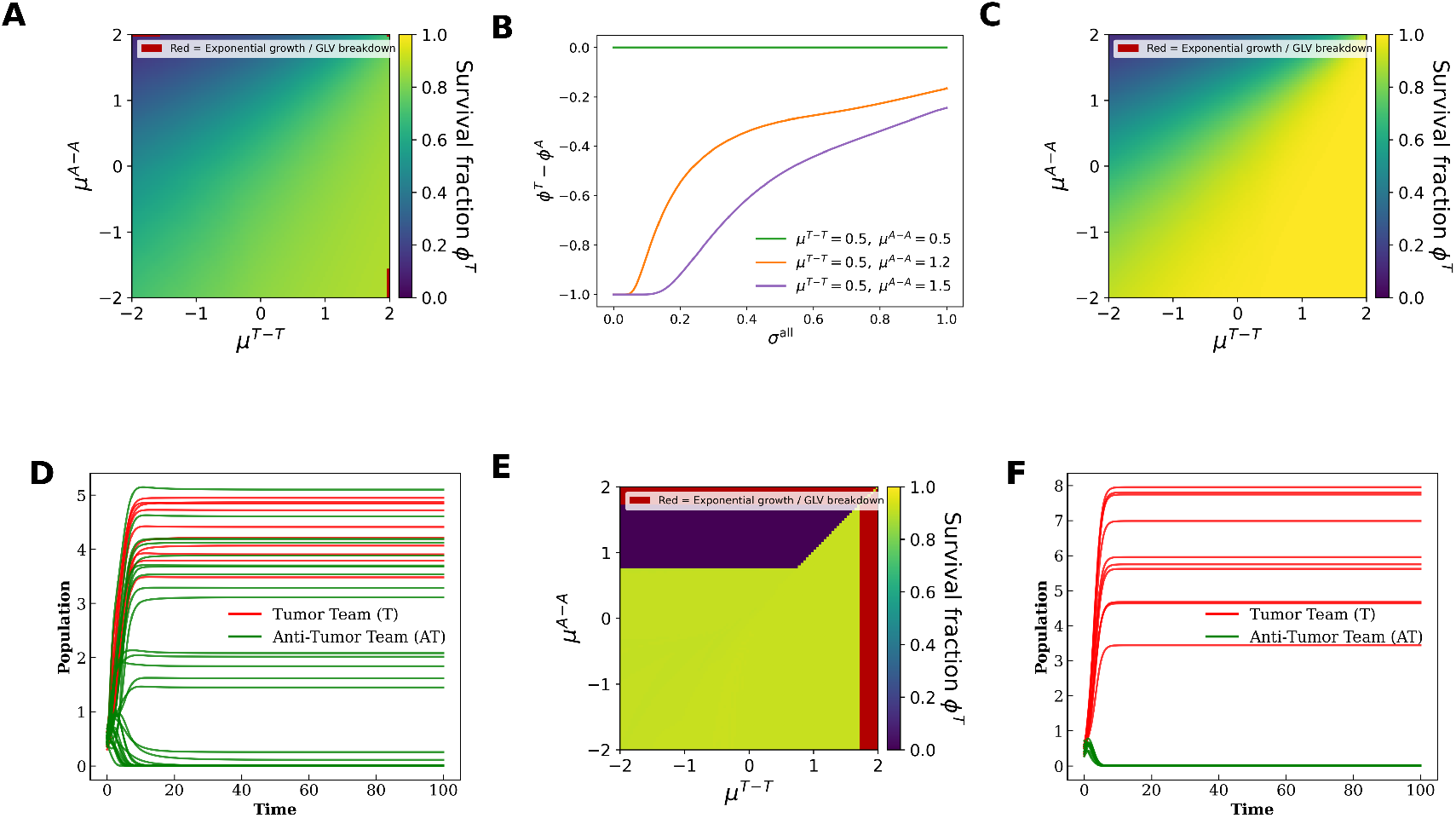
Intermediate heterogeneity leads to extinctions and diffused regimes. (A) Tumor-team survival phase plot when *σ*_all_ = 0.8. (B) Difference in survival fraction between tumor team and anti-tumor team as heterogeneity (*σ*_all_) increases. (E) Tumor-team survival phase plot when *σ*_intra_ = 0.8 and *σ*_inter_ = 0. (C) Survival phase plot when *σ*_inter_ = 0.8 and *σ*_intra_ = 0. *σ*_inter_ is the standard deviation of interactions between the two teams, and *σ*_intra_ is the standard deviation of interactions within the teams. (E) Trajectories of GLVs simulated with 10 members in each team (*σ*_all_ = 0.8, *µ*^*T* −*T*^ = 0.5, *µ*^*A*−*A*^ = 0.5). (F) Trajectories when *σ*^intra^ = 0.8, *µ*^*T* −*T*^ = 1.8, *µ*^*A*−*A*^ = 0.5.

**Figure 7:**
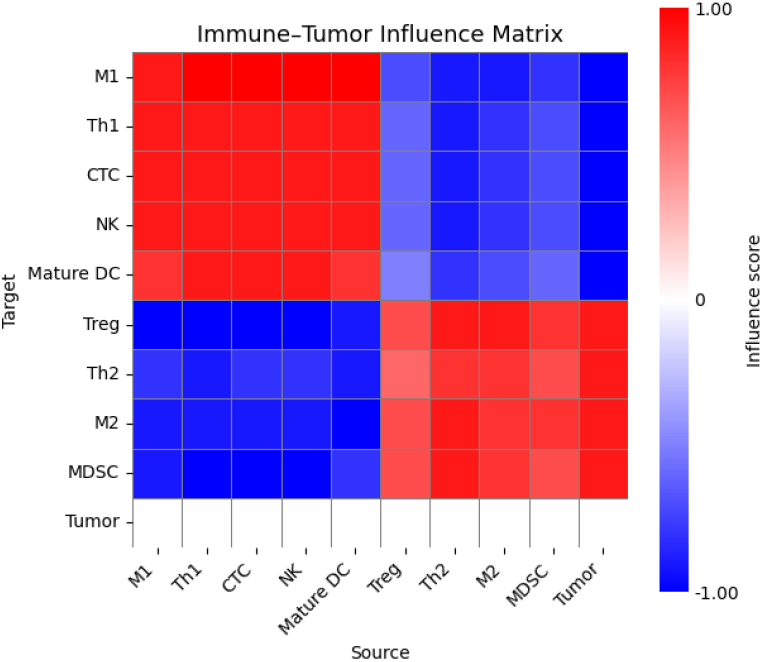
Influence matrix of the interaction network in Figure 1 reveals a strong presence of two teams.

Interestingly, although higher heterogeneity increases the overall likelihood of species loss, it also promotes coexistence. As shown in Fig. 6B, the difference between pro-tumor and anti–tumor survival fractions decreases as *σ*^all^ increases, even though both teams experience reduced survival fractions overall. In other words, heterogeneity leads to coexistence.

Up to this point we used a single heterogeneity parameter,

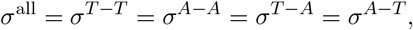

assigning equal variability to all interaction types. We next asked whether specific structural components of the two-team ecosystem are more sensitive to heterogeneity. To address this, we separated variability into two contributions:

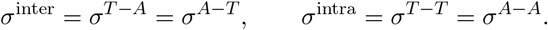

We find that high inter–team heterogeneity produces the strongest coexistence-promoting effect. Variability in cross-team antagonism diffuses phase boundaries and expands coexistence regions across the (*µ*^*T* −*T*^, *µ*^*A*−*A*^) plane (Fig. 6C,D), consistent with our earlier observations for *σ*^all^. In contrast, high intra–team heterogeneity drives the system toward global extinctions without promoting coexistence: although phase boundaries shift, they remain sharp, and no diffused coexistence region emerges (Fig. 6E). These analytical distinctions are directly reflected in full GLV simulations of two-team systems (Fig. 6F).

## 3 Discussion

The ecological nature of the tumor microenvironment (TME) is increasingly recognized as central to understanding tumor progression and therapeutic response. Within this complex ecosystem, protumor and anti-tumor factors interact through webs of cooperation and antagonism that ultimately shape disease outcomes. While there have been several mathematical models of tumor–immune interactions using a variety of approaches [35], earlier ecological modeling efforts [36, 37] have generally focused on only a few interacting components [38] and relied on rather arbitrary parameter choices. Consequently, a community-level understanding of the TME, in which many functionally distinct immune and stromal populations interact simultaneously, remains largely unexplored.

Ecological communities in natural habitats form large and intricate networks of dynamical interdependencies among their constituent species. Remarkably, recent theoretical work has shown that the behavior of such high-dimensional ecosystems can often be captured by a small number of statistical parameters that describe the distribution and structure of interspecies interactions [34]. This insight provides a powerful framework for studying ecological dynamics without requiring detailed knowledge of every individual interaction. We follow a similar approach here and model tumor immune interactions as a two-team ecosystem driven by generalized Lotka-Volterra equations where the interactions between cell types are described statistically. Our coarse-grained representation allows us to move beyond pairwise or low-dimensional models and to study instead how collective behaviors emerge from the interplay of cooperation within teams and competition between them.

Our ecological framework reveals a clear structure in how cooperation within the pro-tumor and anti-tumor teams govern their coexistence. When intra-team cooperation in both communities remains below a certain level, the two teams can coexist, reflecting an early immune-controlled microenvironment where neither side overwhelmingly dominates. However, when cooperation within one team exceeds this threshold, the system undergoes competitive exclusion: the team with stronger internal support suppresses the other. If cooperation becomes high in the pro-tumor community, the anti-tumor team is driven to extinction, a scenario reminiscent of late tumor progression, when tumor-promoting signaling overwhelms local immune control. Together, these results highlight how shifts in internal cooperation within each community can tip the balance between coexistence, immune control, and tumor dominance.

In our model, interaction parameters within and between teams are drawn from Gaussian distributions, and we assume that the corresponding fluctuations are independent. While earlier work on random ecosystems introduces a correlation parameter *γ* between interaction fluctuations, we find that our main results are insensitive to the choice of *γ* (Supplementary Figure X, and a general derivation in presented in the Appendix). The associated standard deviations quantify ecological heterogeneity. Such heterogeneity has been shown to play a central role in tumor progression [6] and is known to modulate tumor–immune dynamics [39]. . Within this framework, we find that heterogeneity shapes ecological outcomes between pro-tumor and anti-tumor factors and can promote coexistence. Notably, the effects of intra-team and inter-team heterogeneity are not symmetric: while increasing intra-team variability generally destabilizes coexistence, inter-team heterogeneity robustly expands the parameter regimes in which both teams persist.

Ratios of pro-tumor and anti-tumor cell populations are widely used as prognostic markers, for example the M1/M2 macrophage ratio [40] and the N1/N2 ratio in tumor-associated neutrophils [21]. The dynamics between these factors individually can be exploited for theraputic benefit [41]. Our model highlights the role of heterogeneity, and in particular distinguishes between intra-team and inter-team heterogeneity as an independent dimension shaping the ecological dynamics. Moreover, recent work has demonstrated that ecological interaction parameters can be inferred, at the level of large ecosystems, from multiplexed imaging datasets of the tumor microenvironment, and that heterogeneity in these parameters can be quantitatively characterized [42]. Our framework can therefore be used to study ecosystem-level dynamics of the tumor–immune microenvironment informed directly by experimentally inferred interaction statistics.

The idea of “teams” as functional modules appears in several biological contexts. For example, the gene regulatory network underlying Epithelial-Mesenchymal Plasticity (EMP) consists of two antagonistic teams of genes whose internal cooperation and mutual inhibition generate rich decision-making dynamics [43]. The relevance and emergent behavior of such team-like architectures have been highlighted across diverse biological systems [44, 45]. Related work has also examined the dynamics of multi-team networks in gene regulatory systems [46]. Extending this framework to ecological settings presents a natural next step, allowing us to investigate how multi-team interactions shape community structure and stability in more complex ecosystems.

A key feature of the tumor microenvironment that we have not incorporated is phenotypic plasticity. Tumor cells, for instance, can reprogram macrophages from an anti-tumor (M1) phenotype to a pro-tumor (M2) state [19], effectively shifting individuals from the anti-tumor team to the pro-tumor team. Similar plasticity has been observed in tumor-associated neutrophils, where N1 neutrophils can transition into N2 neutrophils in response to tumor-derived signals [20].

Such transitions between functional groups are not captured by classical generalized Lotka– Volterra dynamics, which assume fixed species identities and interaction coefficients and do not allow for phenotypic transitions. Extending the present framework to incorporate phenotypic plasticity, as well as interaction coefficients that depend on tumor burden, represents an important direction for future work. Such extensions could reveal how plasticity between the two teams reshapes coexistence, competitive exclusion, and ecosystem-level outcomes. Moreover, tumor-burden-dependent interaction parameters may uncover rich temporal dynamics in the tumor–immune ecosystem.

## Acknowledgments

VA and HL acknowledge support by NSF-PHY2019745. MKJ was supported by Param Hansa Philanthropies.

## Data and Code Availability

Solver used for the self consistent equations and code for generating the figures are available at https://github.com/anandwhybhav/tumor_anti-tumor_teams/

## Author Contributions

HL and VA designed the project, VA did the calculations, and HL, VA and MKJ wrote the manuscript.

## 4 Methods

### 4.1 Simulations

The **interaction matrix** *A* of size *S* = *n*_*T*_ + *n*_*A*_ = 20 was drawn from a Gaussian ensemble 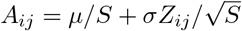, with corr(*A*_*ij*_, *A*_*ji*_) = *γ* and zero diagonals.

**GLV dynamics** followed 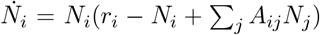 with *r*_*i*_ ∼ 𝒩 (1, 0.0), integrated using <monospace>solve ivp </monospace>(RK45, tol. 10^−6^/10^−8^) to *T* = 300. Agreement between simulations and theory was quantified via *R*^2^ (<monospace>scikit-learn</monospace>) and visualized in <monospace>Matplotlib 3.8</monospace>.

### 4.2 Cavity Solution

A detailed derivation is presented in the Appendix. For the random ecology, We solve the self-consistent equations using a solver implemented in Python. We implemented a self-consistent **cavity method solver** for the two-team generalized Lotka–Volterra (GLV) model, iterating truncated Gaussian moment equations for survival fractions (*ϕ*^*T*^, *ϕ*^*AT*^) and abundance moments 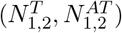 until Δ *<* 10^−8^ or 1000 iterations (relaxation *α* = 0.4).

## 5 Appendix

### 5.1 Influence Matrix

To capture multi-step interactions in the network, we computed an influence matrix from the adjacency matrix *A*. We first generated a binary matrix *A*_max_ by setting all nonzero entries of *A* to one. For each path length ℓ = 1, …, 10, we computed the weighted walk matrix *A*^ℓ^ and the corresponding unweighted walk count 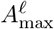. The influence contributed by paths of length ℓ was defined element-wise as

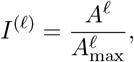

with entries set to zero where 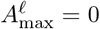. The final influence matrix was obtained by averaging over all path lengths:

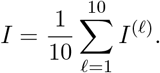

This provides a normalized measure of indirect influence mediated via feedbacks loops of up to length 10.

### 5.2 Generalized Lotka-Volterra (GLV) Framework

The standard form of the generalized Lotka–Volterra (GLV) equations is given by

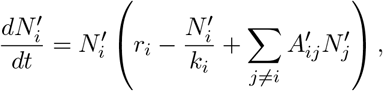

where 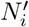 is the abundance of species *i, r*_*i*_ its intrinsic growth rate, *k*_*i*_ its carrying capacity, and 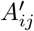 the interaction strength between species *i* and *j*.

We define a rescaled variable 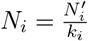, which yields

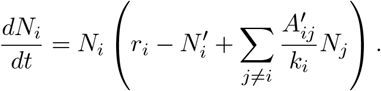

Defining 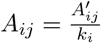, we obtain

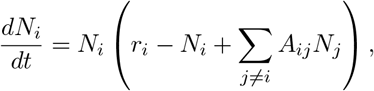

with interactions drawn from a random ensemble,

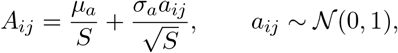

where *S* is the total number of species. In this formulation, the carrying capacity is absorbed into *N*_*i*_ and *A*_*ij*_; thus, *N*_*i*_ *>* 1 represents population levels exceeding the nominal carrying capacity, indicating a cooperative regime.

—

### 5.3 Cavity Derivation with Interaction Reciprocity

Following Bunin [29] and Barbier [30], we consider a generalized Lotka–Volterra (GLV) ecosystem with random interactions. The interaction matrix is written as

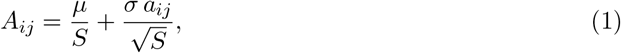

where *S* is the total number of species, *a*_*ij*_ are zero-mean, unit-variance random variables, and correlations between reciprocal interactions are controlled by

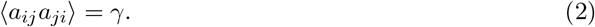

We extend this framework to an ecosystem composed of multiple groups (or “teams”) indexed by *x* ∈ {1, 2, … }. The abundance 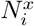 of species *i* in group *x* obeys

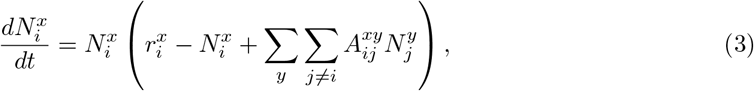

with interaction coefficients

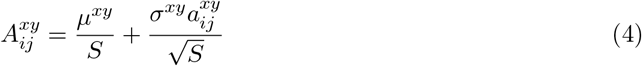

We denote by *S*^*x*^ the number of species in group *x*, and define the group population means

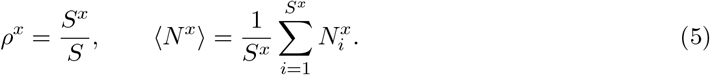

Separating deterministic and fluctuating contributions in Eq. (7), the interaction term may be approximated as

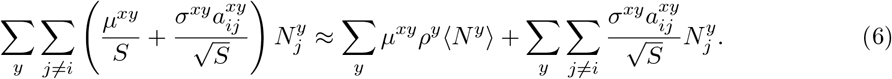

Substituting the above expression in the GLV results in

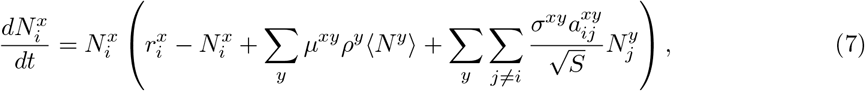

We absorb the deterministic term ∑_*y*_ *µ*^*xy*^*ρ*^*y*^⟨*N*^*y*^⟩ into an effective growth rate defined as

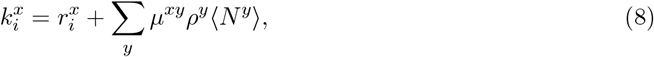

the dynamics reduce to

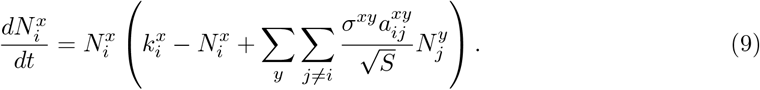

The cavity method proceeds by introducing an additional cavity species that perturbs the ecosystem weakly through its interactions with the resident species. The response of the system feeds back onto the cavity species via these interaction parameters, allowing the dynamics of the cavity species to be expressed in terms of the collective state of the system. The central cavity approximation assumes that, due to the generic and statistically homogeneous nature of the interactions, the effective dynamics experienced by the cavity species is representative of the dynamics of a typical species in the full system. This assumption leads to a closed set of self-consistent equations for parameters, which are solved numerically.

We now introduce a cavity species 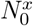 in group *x*, whose dynamics obey

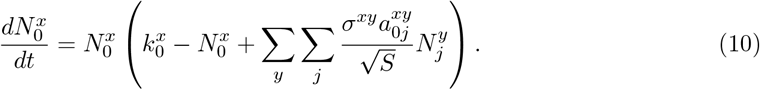

The presence of the cavity species perturbs the resident ecosystem through shifts in the effective growth rates

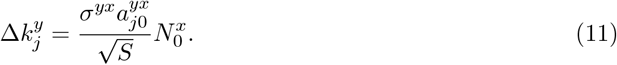

We assume that each species in the ecosystem responds linearly to perturbations in the effective growth rates due to the cavity species and we get.

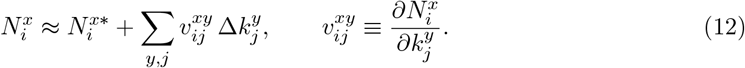

Where 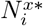 is the unperturbed equilibrium abundance and 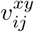 is the response coefficient of *i*^*′*^*th* species in group *x* due to growth rate perturbations in *j*^*′*^*th* species in group *y* Substituting this response into the fluctuating term of GLV equation for the cavity species (10) yields

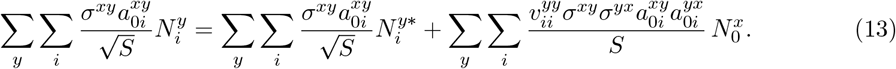

From the above expression, we define

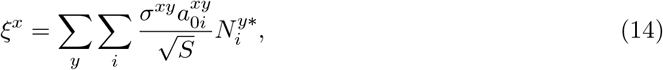

Substituting the reciprocity parameters, 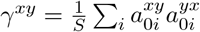 we obtain

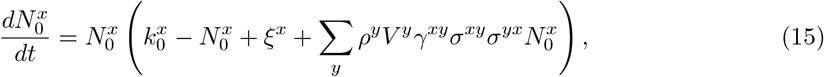

where 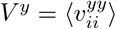.

At equilibrium 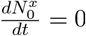, and for 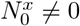 we have,

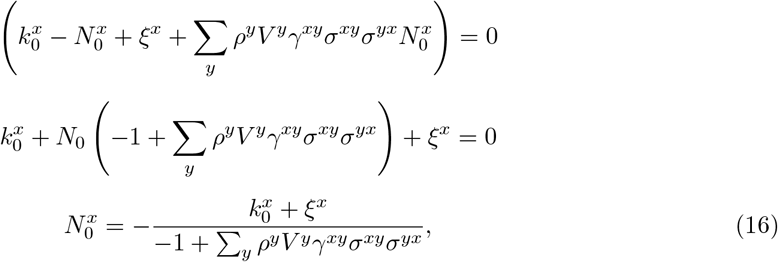

We define 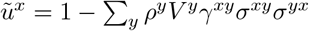. That results in the expression

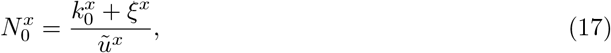

Since abundances must be non-negative, the solution is truncated at zero:

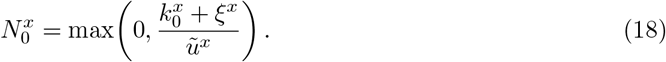

The deterministic and stochastic components are distributed as

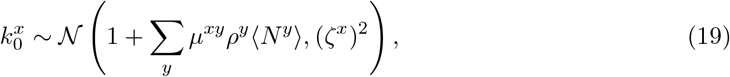

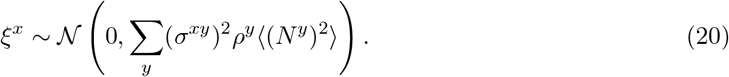

Thus, the distribution 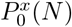 is a truncated Gaussian with mean and variance

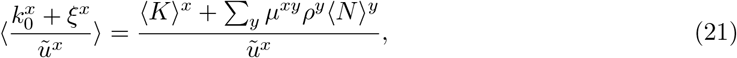

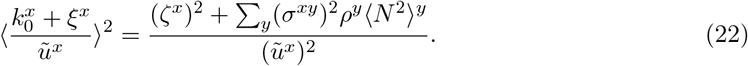

Let’s use 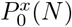 to denote the Gaussian distribution induced by 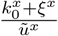 then *N*_0_ is the distribution restricted to positive values giving us the following self consistent equations

#### 5.4 Self-Consistent Equations

For each group *x*, the parameters satisfy

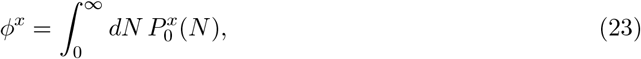

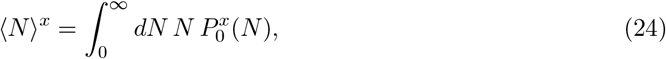

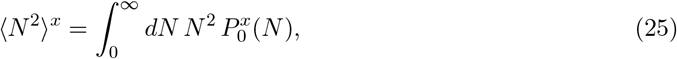

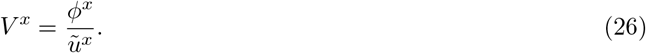

#### 5.5 Two-Team Structure

To incorporate a “two-team” ecological structure, we define *x, y* ∈ {*T*, A} representing tumor-promoting and anti-tumor groups, respectively. Interaction means are assigned as:

- *µ*^*T* −*T*^, *µ*^*A*−*A*^ *>* 0 for intra-team cooperation (self-activation),
- *µ*^*A*−*T*^, *µ*^*T* −*A*^ *<* 0 for inter-team antagonism (*x* ≠ *y*).

The parameter *ρ*^*T/A*^ quantifies the relative fraction of species within team *T/A*. These coupled self-consistent equations govern the steady-state statistics and coexistence regimes emerging from the competition and cooperation between the two teams.

